# Characterization of aquaporin1b (AQP1b) mRNA in mud loach (*Misgurnus mizolepis*) in response to heavy metal and immunostimulant stimuli

**DOI:** 10.1101/2021.09.09.459705

**Authors:** Sang Yoon Lee, Yoon Kwon Nam, Yi Kyung Kim

## Abstract

Aquaporins (AQPs) facilitate the transport of water or other small solutes into cells in the presence of osmotic gradients. However, the current understanding of piscine AQP gene with cellular stress responses has been still limitedly exemplified. In present study, we characterized the mud loach AQP1b gene at the nucleotide and amino acid levels. We identified three AQP 1b transcript variants (mmAQP1b_tv1, mmAQP1b_tv2, and mmAQP1b_tv3). Then, we examined the AQP1b promoter region and observed several transcription factor binding sites (TFBS) for nuclear factor of activated T-cells (NFAT), SRY-box, c-AMP responsive element binding protein (CREB), GATA binding factor, and hepatic nuclear factor-1. Interestingly, mmAQP1b transcription was differentially modulated by heavy metal or immunostimulant challenge. Further studies to deepen the knowledge of fish AQP-mediated adaptation response potentially relevant to molecular pathogenesis are warranted.

**Summary statement:** We identified mud loach AQP1b transcript variants and consensus sequences involved in stress or innate immunity in promotor region. AQP1b transcription was differentially modulated by heavy metal or immunostimulant challenge.

## Introduction

Teleost species have the remarkable ability to withstand acute or long-term fluctuations in environmental salinity. Fish cells monitor the osmolality difference between the intracellular and extracellular spaces, and transport water to recover their volume following cell swelling or shrinkage. Salinity stress is linked to a wide range of biological processes *e.g.* metabolism, mortality, growth, and even immune responses (Baltzegar et al., 2014; Moshtaghi et al., 2016). After being subjected to an imbalance between environments, proteins trigger various complex responses such as changes in structure or function, a consequence of which is altered enzyme activity (Fiol and Kültz, 2007).

Aquaporins (AQPs) facilitate the transport of water or other small solutes into and out of cells in the presence of osmotic gradients. These integral membrane proteins have been identified across phyla, from Archaea (Kozono et al., 2003) to primates (King et al., 2004). Based on amino acid sequence similarity, 13 different AQPs (AQPs 0-12) can be divided into three subfamilies: classical aquaporins that selectively transport water; aquaglyceroporin that transport glycerol and other small molecules in addition to water; and an unorthodox subgroup (Zardoya, 2005; Ishibashi et al., 2011). Genomic and phylogenetic analyses revealed that teleosts possess several AQP isoforms which have undergone lineage-specific changes and divergence via whole-genome duplication events (Tingaud-Sequeira et al., 2010). Zebrafish (*Danio rerio*) shows a much higher diversity of AQPs than tetrapod AQPs. Duplicated or triplicated sub-isoforms have been retained in the zebrafish genome (Tingaud-Sequeira et al., 2010; Finn and Cerdà, 2011; Finn et al., 2014; Madsen et al., 2015). The valuable information regarding the entire aquaporin sequence in water fleas (*Daphnia pulex*) have been recently published in the NCBI non-redundant protein database (Lind et al., 2017).

Many studies indicate that AQPs show multiple modes of activation and regulation, enabling them to respond to diverse cellular events such as neutral signal transduction, brain swelling, and cellular migration (Balzegar et al., 2014; Moshtaghi et al. 2016). To date, understanding the physiological role of piscine AQPs has been facilitated by studying the expression of AQPs in response to changes in salinity (Cutler and Cramb, 2002; Watanabe et al., 2005; Giffard-Mena et al., 2011; Kim et al., 2010; Choi et al., 2013; Madsen et al., 2015).

Mud loaches (*Misgurnus mizolepis*; Teleostei; Cypriniformes) are currently the most popular freshwater species with growing importance of domestic market in Korea. In addition to its commercial importance, the mud loach has attractive merits as an experimental organism, including small adult size, high fecundity, year-round spawning under controlled conditions, and relatively well-established techniques for genetic manipulation (Nam et al., 2011; Cho et al., 2012). Given these advantages, mud loaches could be particularly relevant for studying the involvement of aquaporins in physiological processes.

Most studies investigating piscine aquaporin genes focus on the effects of salinity-induced adaptation in piscine AQP genes in response to biological challenges. However, investigating the interaction among genes using molecular genetic approaches has elucidated the molecular mechanisms underlying biological function and regulatory gene expression. Identifying interaction among diverse genes and autonomous adaptation effects are essential for systematically unraveling the cellular or molecular mechanisms of AQP action. Recent reports indicate that AQPs could be involved in inflammasome activation-induced cell volume regulation (Compan et al., 2011; Meli et al., 2018). Previously, we identified a novel AQP1a in mud loaches (mmAQP1a) which is differentially modulated *in vivo* (Lee et al., 2017). In fact, the existence of AQP1a and its duplicate in teleost species is well established, showing specialized expression (Zapater et al., 2011). We found a AQP1a paralog from our next-generation sequencing database for the mud loach (unpublished data). The genetic determinants for the stenohaline freshwater species mud loach AQP1a paralog, called AQP1b have not been extensively explored. In present study, we identified the AQP1a paralog in mud loaches. The AQP1a paralog, AQP1b is a teleost-specific water-selective channel that mediates oocyte hydration, which is based on the coordinated action between osmolyte flux and aquaporin activity (Fabra et al., 2005; Sun et al., 2010). We found three variant AQP1b paralogs that are structurally similar to mmAQP1a. In line with our long-term goal to more deeply understand the involvement of aquaporin molecules in osmoregulation and physiological processes of mud loaches, we aimed to identify the mRNA splice variants of AQP1 in mud loach. We characterized the structural and features of the AQP1 promoter and examined gene expression patterns in response to environmental challenges.

## Materials and methods

### Isolation of AQP1b cDNA from mud loaches

To isolate the aquaporin cDNA sequence, a mud loach expressed sequence tag (EST) database generated from whole body RNAs was surveyed. Based on the EST survey and a homology search of NCBI GenBank (http://blast.ncbi.nlm.nih.gov/), the partial mud loach AQP sequences were identified. The full-length of mmAQP sequence was obtained from mud loach total RNAs by RT-PCR and/or Vectorette PCR isolation using specific primer pairs (Table 1). At least six clones for each AQP isoform were sequenced to obtain the representative sequence.

### Cloning the mud loach AQP1b gene and promoter

Based on the full-length cDNA sequences of the mud loach AQP, the gene sequence was isolated using PCR with AQP1-specific primers that bound the untranslated region (UTR) of gene. The oligonucleotide primer pairs and PCR conditions used are shown in Table 1. The amplified PCR products were cloned into pGEM-T easy vector (Promega, Madison, WI, USA). Sequencing for recombinant clones (n = 6) that contained the correct insert size were done by primer walking with gene-specific primers. To isolate the 5’-flanking region, genome walking was performed using a Universal Genome Walker Kit (Clontech Laboratories Inc., USA). The DNA fragments obtained from genome walking were subcloned, sequenced, and assembled into a contig as described. Finally, the AQP genomic DNA spanning the isolated region was PCR amplified. The PCR products were sequenced to obtain the representative genomic sequence of the mud loach AQP gene.

### Bioinformatic sequence analysis

The cDNA sequence was analyzed using the Open Reading Frame (ORF) Finder tool (https://www.ncbi.nlm.nih.gov/orffinder/). Using the deduced amino acid sequence, the theoretical molecular weight and isoelectric point (pI) were evaluated with the ExPASy ProtParam online tool (http://web.expasy.org/protparam/). A homology search of the mud loach AQP1 was performed using the NCBI BLASTx algorithm (https://blast.ncbi.nlm.nih.gov/Blast.cgi) and/or the Ensembl genome database (http://asia.ensembl.org/index.html). Multiple sequence alignment of the aquaporin amino acid sequence to representative teleostean and human orthologs (Table 2) was performed using CLUSTAL W (Thompson et al., 1994). Transmembrane domain prediction was performed using the TMHMM Server v. 2.0 (http://www.cbs.dtu.dk/services/TMHMM-2.0/). To gain insight into the transcriptional regulation of AQP, putative transcription factor binding sites (TFBSs) in the proximal promoter region were identified using MatInspecter software (http://www.genomatix.de) This prediction was performed using the default parameters [General Core Promoter Elements (0.9 core / Optimized matrix sim) and Vertebrates (0.9 core / Optimized matrix sim)] based on Matrix Family Library Version 11.0 (September 2017). We then characterized the regulatory regions of AQP1a.2 gene promoter by sequencing 3405 bp from the 5′-flanking region and analyzed this region for *cis-*regulatory elements using MatInspector software.

### Tissue sampling for basal expression assays

To investigate the constitutive expression of AQP in various adult mud loach tissues, we collected samples from 13 different tissues (brain, eye, fin, gill, heart, intestine, kidney, liver, muscle, skin, spleen, ovary, and testis) w from six-month-old healthy females (mean body mass = 19 ± 4 g; *n* = 8 for each group) and males (mean body mass = 12 ± 3 g; *n* = 8 for each group). The tissues from the sampled individuals were pooled prior to total RNA extraction.

### Sampling for embryos and early larvae

Fertilized eggs were collected by mating female fish with male fish using the “wet method” (Nam et al., 2004). The fertilized eggs [referred to as zero hours post-fertilization (HPF); hour 0] were incubated and kept at 25 ± 0.5°C and under an ambient photoperiod until hatching. The embryonic developmental stages were determined according to Nam et al. (2004). The embryos were pooled at the 32 cell (2 HPF), early blastula (4 HPF), early gastrula (6 HPF), late gastrula (8 HPF), 3-4 myotome (12 HPF), 16-17 myotome (16 HPF), and 23-24 myotome (20 HPF) stages. Hatching stared at 24 HPF and 100% of the embryos hatched by 28 HPF. After hatching, the larvae were transferred to a 50-L tank held at 24 ± 0.5°C. 100 to 150 larvae were collected at 1, 2, 3, 4, 5, 6, 7, and 14 days post-hatching (DPH). Larva rearing was performed as previously described (Nam et al., 2004).

### In vivo stimulatory treatments

We examined AQP1b mRNA expression following acute exposure to cadmium (Cd), chromium (Cr), copper (Cu), iron (Fe), manganese (Mn), nickel (Ni), or zinc (Zn). Fish (n = 12, average body mass = 15.2 ± 4.1 g) were separated into seven experimental groups (one per metal) and one control group and kept in 60 L tanks with the respective treatment. The tanks were filled with tap water and held at 25 ± 0.5°C. Each group was acclimated to the tanks for one week. The dose for each metal was 5 μM (Cho et al., 2009), and the exposure time was 48 h. All chemicals were of analytical or reagent grade and were obtained from Sigma-Aldrich. No feed was supplied during the exposure period. The tank water was changed every day to maintain the desired metal concentration. The control group was kept in the same condition as the other groups without the addition of any metals. After 48 h of exposure, four tissues (liver, intestine, kidney, and spleen) were collected.

To examine potential immune responses, fish (15.2 ± 4.1 g; *n* = 3 for each group) were intraperitoneally injected with lipopolysaccharide (LPS; *Escherichia coli* 0111:B4, Sigma-Aldrich, St Louis, MO, USA), polyinosinic:polycytidylic acid [poly(I:C); Sigma-Aldrich], or phosphate buffer saline (PBS, pH 6.8; non-stimulated control). For LPS or poly(I:C) challenge, two doses (5 or 25 μg g^−1^ body weight) were tested. After being injected, each group was transferred to a 60 L tank at 25°C. No feed was supplied during this period. At 24 h post-injection, immune-relevant tissues were surgically removed from each fish and stored at −80°C until analyzed by RT-PCR.

### RT-PCR analysis

Total RNA was extracted using TriPure Reagent (Roche Applied Science, Mannheim, Germany) and purified using an RNeasy Mini Plus Kit (Qiagen, Hilden, Germany), including DNase I treatment. For individual experiments, 2 μg total RNA was used for cDNA synthesis using an Omniscript Reverse Transcription Kit (Qiagen). For the quantitative real-time PCR, the cDNA was diluted 4-fold with sterile distilled water and 2 μL was used as the template for each qPCR reaction. The qRT-PCR analysis was conducted using a LightCycler 480 Real-Time PCR System and LightCycler 480 SYBR Green I Master mix (Roche Applied Science, Germany). 18S RNA (adult tissue and embryonic-early larval samples) and ribosomal protein 7 (RPL7 experimental stimulation analysis) were chosen as the internal control genes. The plasmid DNAs containing the amplified target mRNAs were prepared as standard samples. The qRT-PCR analysis for each experiment included three technical replicates for each biological specimen.

To compare AQP1a.2 transcript levels across tissue types and developmental samples, qPCR data were analyzed using the ΔCt method (Ct of the AQP1b gene subtracted from the Ct of the internal control genes). The relative AQP1a.2 gene expression in the stimulated or challenged groups was analyzed as the fold difference compared to the control group using the 2^−ΔΔCt^ method (Schmittgen and Livak, 2008).

### Statistical analysis

All data are expressed as means ± standard error. The data were analyzed using one-way ANOVA followed by Duncan’s multiple range tests included in SPSS version 10.0 software (SAS Inc., Cary, NC, USA). *P*<0.05 was considered statistically significant.

## Results

### Characteristics of the AQP1b isoform

The AQP1b gene from the mud loach (named mmAQP1b) is 857 bp long with an open reading frame (ORF) encoding 269 amino acids. The calculated molecular mass of this isoform is 28.86 kDa, with a theoretical pI value of 9.35. The nucleotide sequence of the AQP1b cDNA sequence was submitted to GenBank under the accession numbers MT184376-MT184378 (genomic DNA MT184375). The deduced amino acid sequence of mmAQP1b shares the same core architecture as the vertebrate AQPs, including six transmembrane domains and cytoplasmic amino and carboxy termini (Fig. 1). The two hydrophobic loops contain asparagine-proline-alanine (NPA) motifs that regulate selective water conduction and maintain proton gradients across the biological membrane (Gonen and Walz, 2006). The alignment of the putative amino acid sequence of mmAQP1b and orthologous sequences from other teleosts revealed considerable similarity (51-84%) (Table 2). In addition, the cysteine residue near the second NPA motif, which is involved in sensitivity to mercury, was identified in mmAQP1b (Preston et al., 1993).

**Figure 1.**
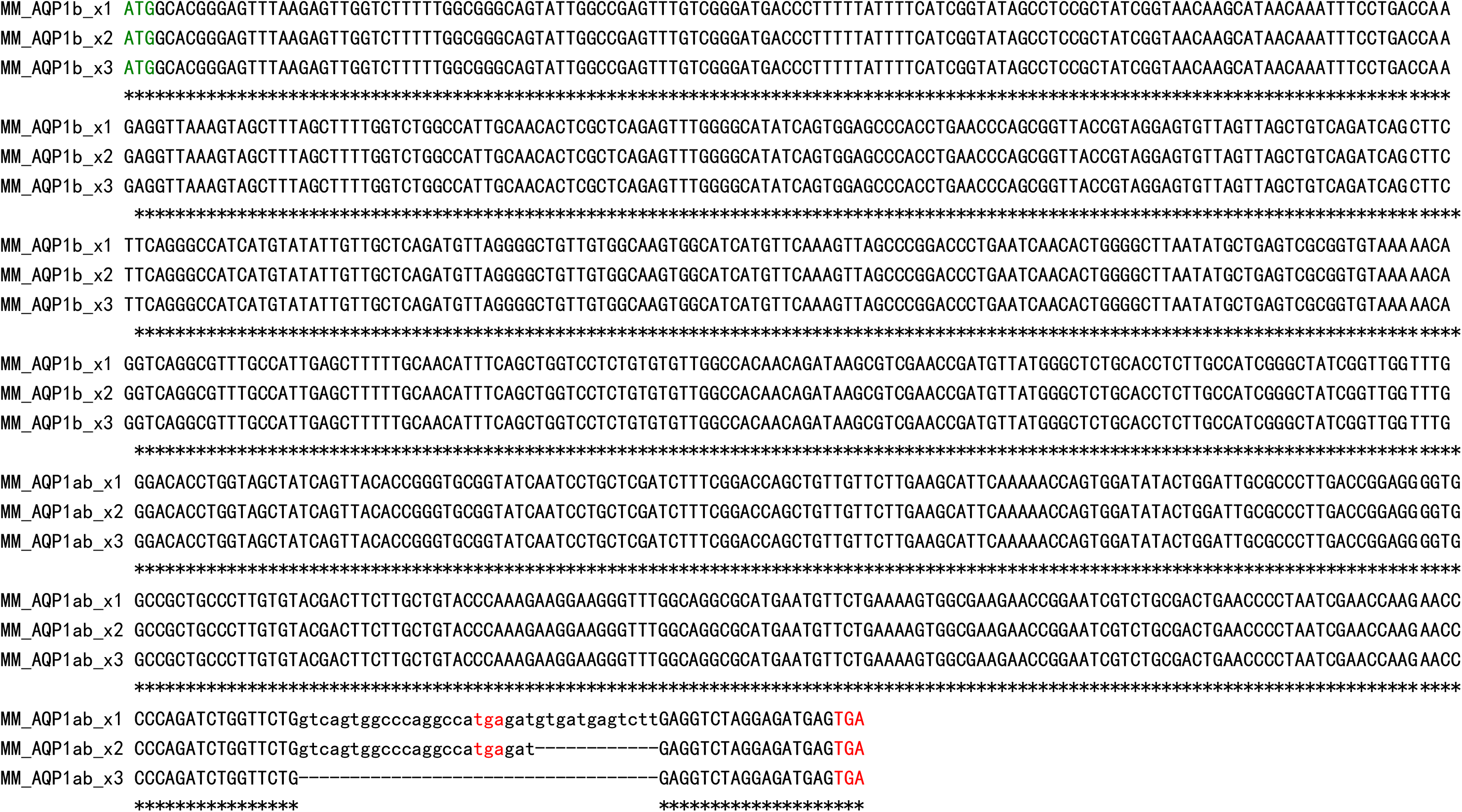
Nucleotide alignment of AQP1b variant tv1, tv2, and tv3 cDNA sequences. Nucleotide gaps are represented by dashes (-).

### Identification of novel splice variants, genomic structure, and organization of mud loach aquaporin 1b

While confirming the full-length mmAQP1b cDNA, we identified three different AQP1b transcript variants, which we designated mmAQP1b_tv1, mmAQP1b_tv2, and mmAQP1b_tv3. These variants were 944, 932, and 819 bp long, respectively. Alignment of the AQP transcripts sequences with other teleosts revealed that the variants arose from alternative splicing between exon 4 and exon 5 for AQP1b_tv2 and AQP1b_tv3. All transcript variants had ATG as the start codon and encoded a 269-amino acids peptide with a molecular mass of 28.9 kDa.

At the genomic level, the mmAQP1b gene contains five exons (366, 165, 81, 178, and 20 bp in length for exon-I to exon-V, respectively) interrupted by four introns (1367, 185, 454, 4036 bp in length for intron-I to intron-IV, respectively). The splice sites contained conserved GT-AG dinucleotides at each junction, and the sequence of each coding exon was clearly matched with its corresponding cDNA sequence.

### In silico analysis of transcription factor binding sites in the mmAQP1b promoter region

To identify the transcription factor regulating mmAQP1b expression, we sequenced a 3405 bp upstream region and analyzed this region for *cis-*regulatory elements using MatInspector software. The transcription start site of AQP1b mRNA was located at −9 bp upstream of the translational ATG initiation codon. We also detected the presence of consensus sequences for core promoter elements important for basal transcription, such as a TATA box (Table 2). In addition, various elements involved with immune modulation or stress responses were observed, including CCAAT-enhancer binding protein (CEBP) sites, cAMP-responsive element binding protein (CREB) sites, nuclear factor of activated T-cells (NFAT) sites, and signal transducer and activator of transcription (STAT) sites. Interestingly, we identified binding sites for three Sry-related high mobility group [HMG]-box (Sox) family members known to be expressed in teleost oocytes. The binding sites for Sox5, Sox6, and Sox3 were located at −2575 bp, −268 bp, and −1048 bp from the transcription initiation site, respectively. In addition, fork head domain factor (Fox), which are involved in development of lung, brain, thymus, and cardiac tissue, had a predicted binding site at −2572 bp based on the consensus sequence. Binding sites for hepatic nuclear factor 1 (HNF1), a transcription factor that regulates ubiquitous expression in many tissues and cells, were also observed at various locations within the AQP1b promoter.

### Tissue distribution and developmental expression of mmAQP1b

mmAQP1b transcript was detected in all examined tissues, although the basal expression level was largely different among these tissues. High AQP1b mRNA expression was detected in the gill, kidney, and spleen, whereas lower expression was observed in the liver, muscle, and skin (Fig. 2).

**Figure 2.**
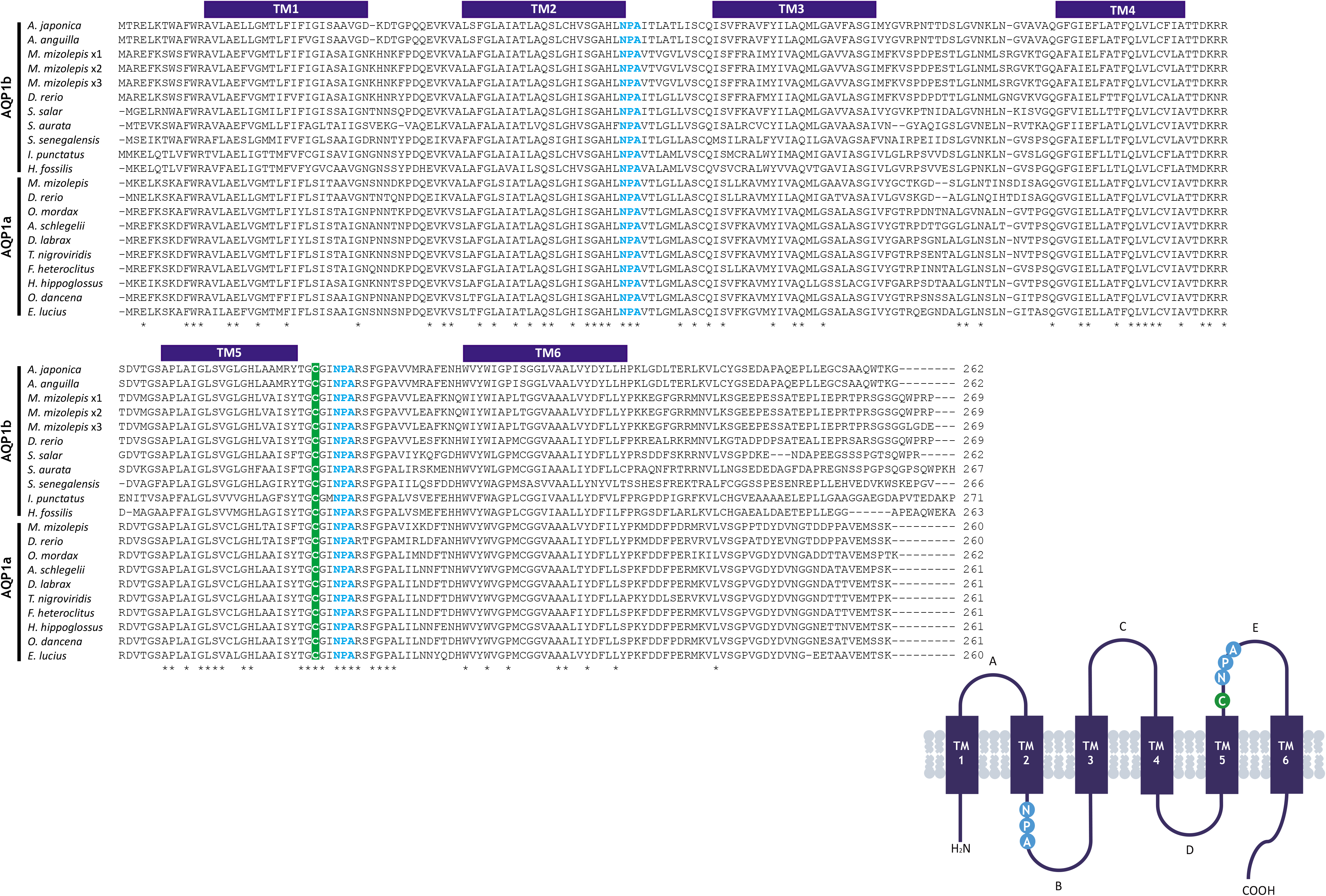
Multiple amino acid sequence alignments and transmembrane topology prediction of AQP1b transcript variants. Asterisks and hyphens indicate identical residues and gaps introduced for optimum alignments, respectively. Two NPA motifs are shown in bold light blue. The locations of 6 putative membrane–spanning domains are shown by navy boxes above the alignment. *A. japonica*: Japanese eel *Anguilla japonica*; *A. anguilla*: European eel *Anguilla anguilla*; *M. mizolepis* tv1, tv2, tv3: mud loach *Misgurnus mizolepis*; *D. rerio*: zebrafish *Danio rerio*; *S. salar*: Atlantic salmon *Salmo salar*; *S. aurata*: gilt-head sea bream *Sparus aurata*; *S. senegalensis*: Senegalese sole *Solea senegalensis*; *I. punctatus*: channel catfish *Ictalurus punctatus*; *H. fossilis*: stinging catfish *Heteropneustes fossilis*; *O. mordax*: rainbow smelt *Osmerus mordax*; *A. schlegelii*: blackhead seabream *Acanthopagrus schlegelii*; *D. labrax*: European bass *Dicentrarchus labrax*; *T. nigroviridis*: green pufferfish *Tetraodon nigroviridis*; *F. heteroclitus*: mummichog *Fundulus heteroclitus*; *H. hippoglossus*: Atlantic halibut *Hippoglossus hippoglossus*; *O. dancena*: marine medaka *Oryzias dancena*; *E. Lucius*: northern pike *Esox Lucius.*

The mRNA expression level of mmAQP1b was regulated during embryonic and larval development (Fig. 3). mmAQP1b expression was low at fertilization (0 hpt) and further decreased until 6 hpt (early gastrula stage). Expression gradually begun to increase with developmental stage until 28 hpt. Afterward, mmAQP1b expression in embryos was remarkably increased until hatching, and dynamically increased even more until 2 dph. Then, mmAQP1b mRNA decreased to the level observed at 28 hpt, although the rate of decrease slowed 4 to 7 dph.

**Figure 3.**
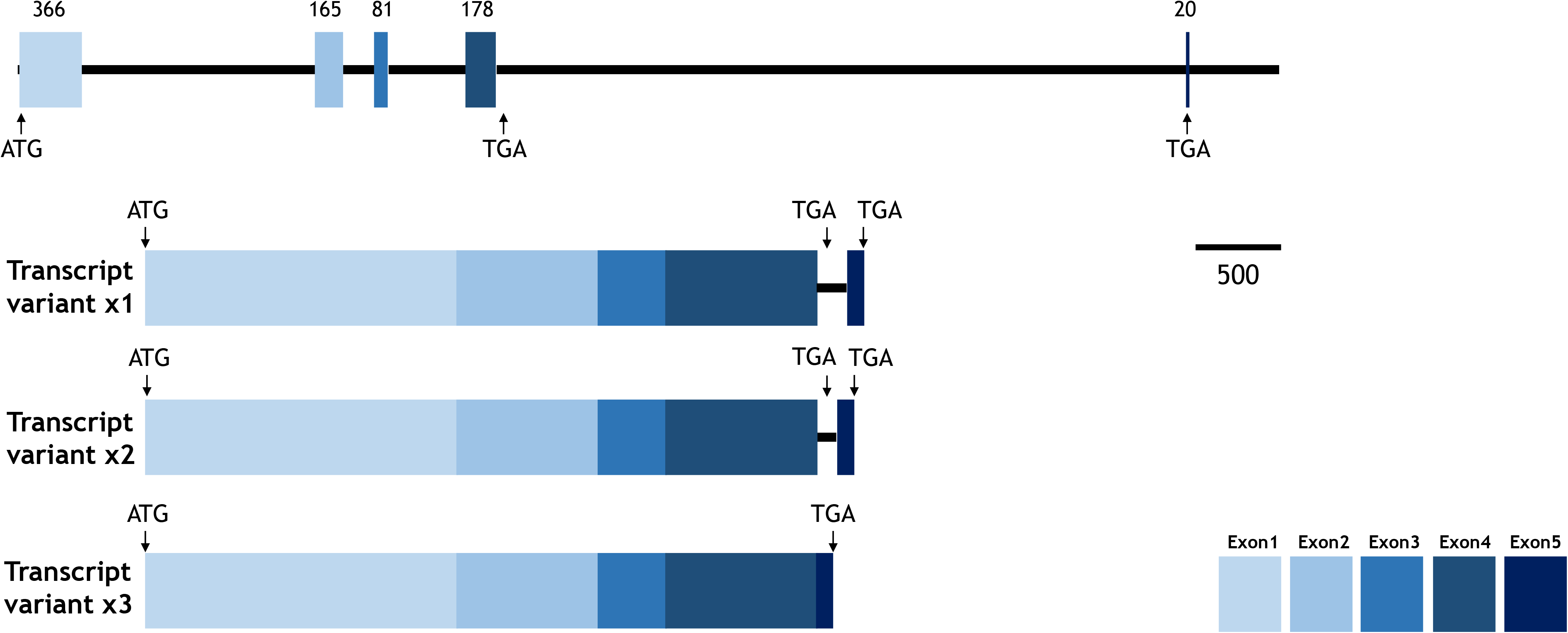
Gene organization and mRNA variants of the aquaporin 1ab gene in mud loaches. mmAQP1b genes contain 5 exons. Each exon is represented by a different color and by Roman numerals. The mRNA transcript length is listed on each exon.

### Modulation of mmAQP1b in response to heavy metal exposure

The response of mmAQP1b to acute waterborne metal exposure was variable according to the metal and tissue type. In the intestine, AQP1b transcription was upregulated by most heavy metals. Cu and Fe caused 3.6- and 2.0-fold decreases of AQP1b transcription, respectively. Ni exposure did not significantly alter mRNA transcription. In the kidney, transcriptional suppression occurred in the groups exposed to Cu (7.3-fold decrease), Fe (3.2-fold), and Mn (2.0-fold). The maximum induction of AQP1b transcript in the kidney was observed with Cr (2.12-fold) treatment. Meanwhile, hepatic AQP1b transcription was induced by all tested metals. Three metals, Cd (25.2-fold increase relative the non-exposed control), Cr (15.1-fold), and Ni (9.18-fold) induced more AQP1b transcription in the liver than the other heavy metals: Cu (5.1-fold) and Zn (2.59-fold). In the spleen, four metal-treated groups did not show increased AQP1b mRNA. Of the four groups, one group (Cu) displayed significantly reduced AQP1b mRNA after challenge. Maximum inducibility in the spleen was 2-fold (the Cu-exposed group), while the other treatments induced only moderately increased AQP1b transcription from 1.19-1.28 folds.

### AQP expression after immune challenge

Experimental challenge with LPS and poly(I:C) altered AQP1b gene expression in many groups, and the patterns were variable among tissues types (Fig. 5). In the intestine, mmAQP1b mRNA was reduced by LPS or poly(I:C) challenge (each 1.27-fold lower than the relative PBS-injected group). In contrast, renal AQP1b expression was significantly upregulated in all challenge groups, with 2.21-fold or 3.34-fold increases relative to PBS-injected controls. A similar pattern was observed in the spleen. AQP1b mRNA expression in the spleen was induced by LPS (2.1-fold) and poly(I:C) (1.93-fold). In the liver, LPS downregulated AQP1b mRNA by 2.32-fold. Other challenges did not show significant differences.

**Figure 4.**
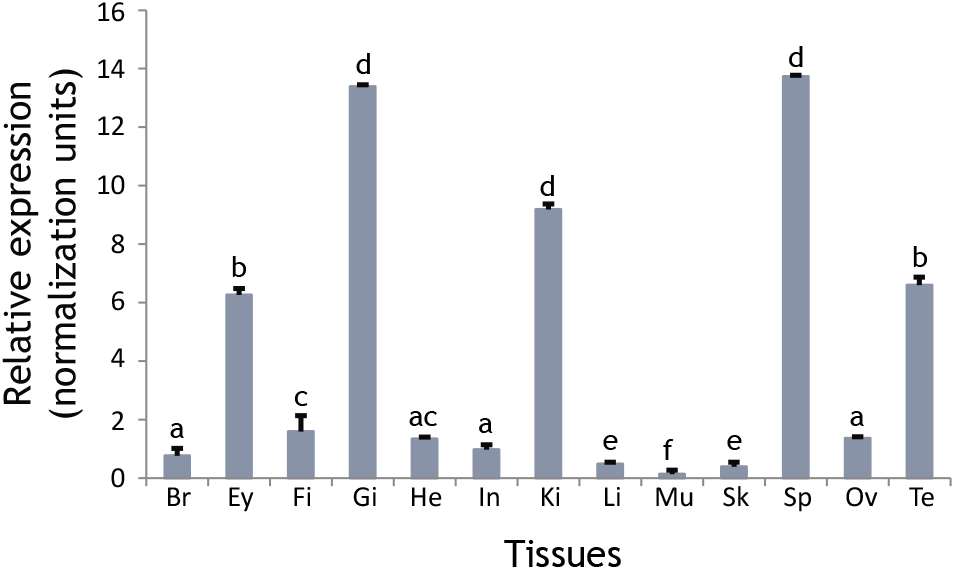
Tissue distribution of mmAQP1b in adult tissues. Abbreviations: brain (Br), eye (Ey), fin (Fi), gill (Gi), heart (He), intestine (In), kidney (Ki), liver (Li), muscle (Mu), spleen (Sp), ovary (Ov), and testis (Te).

**Figure 5.**
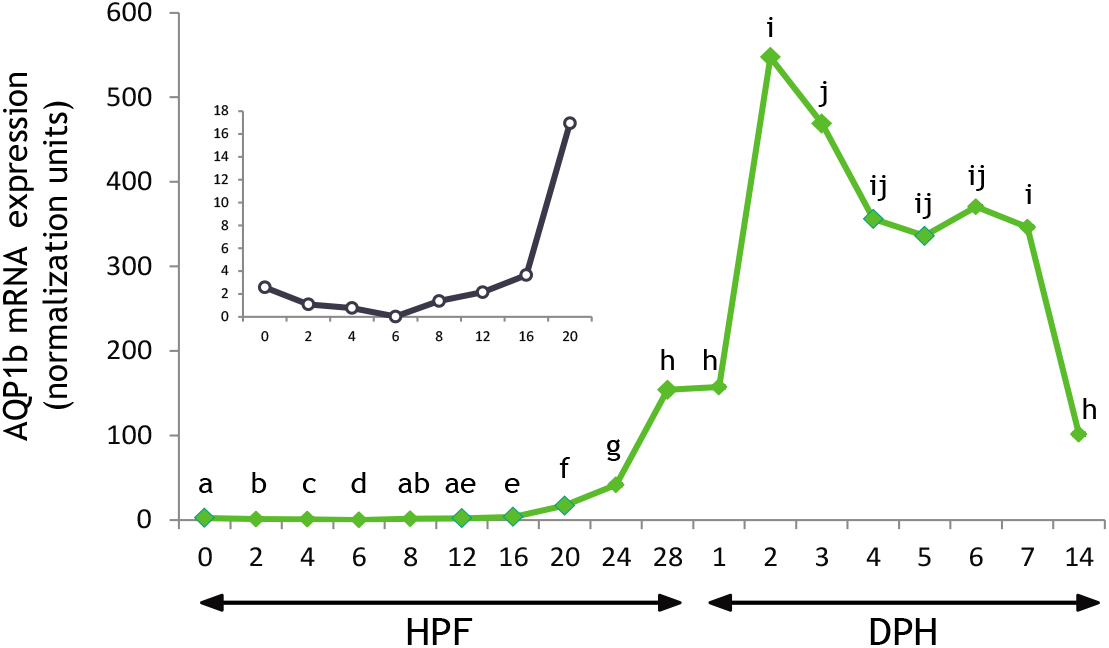
Expression of mmAQP1b mRNA during embryogenesis and larval development. Variant transcripts in different tissues under abiotic stress. Data are represented as means ± SDs. Letters indicate significant difference (one-way ANOVA).

**Figure 6.**
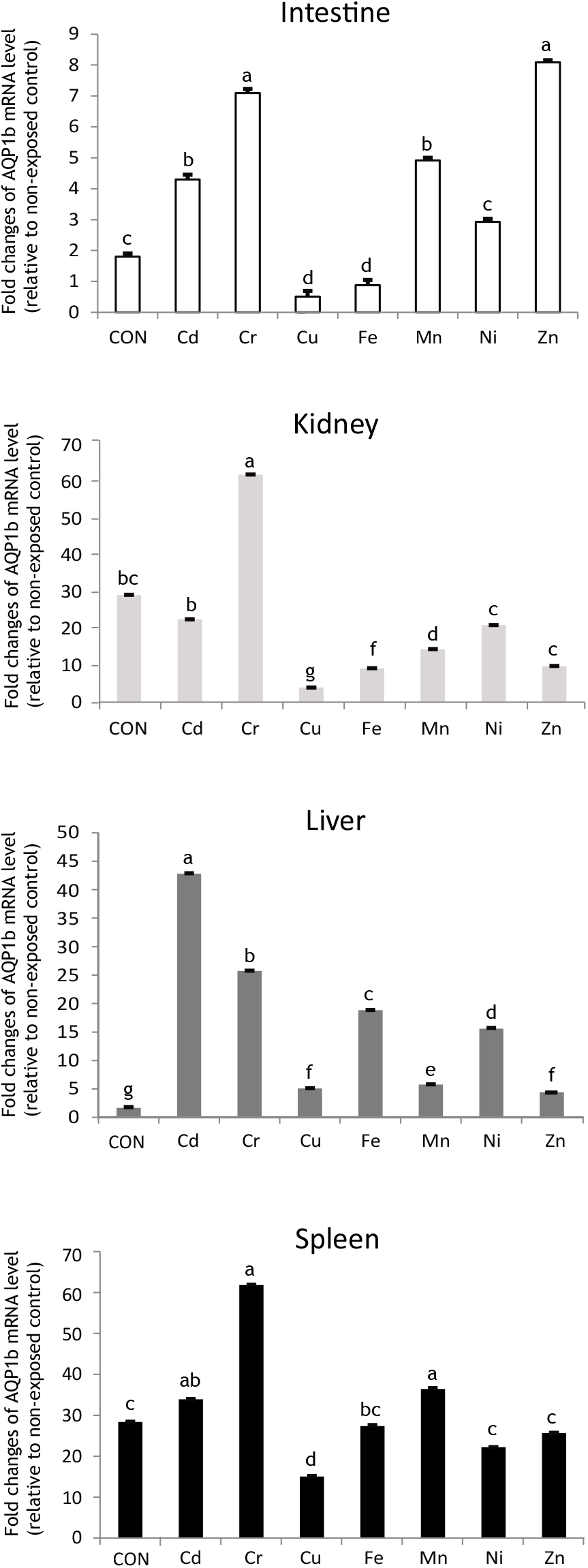
Transcriptional responses of mmAQP1b to acute metal exposure in different tissues. AQP1b expression in metal-exposed groups are expressed as fold changes relative to the non-exposed control group. Data represent means ± SDs. Different letters indicate significant differences, as analyzed by ANOVA followed by Duncan’s multiple range tests.

**Figure 7.**
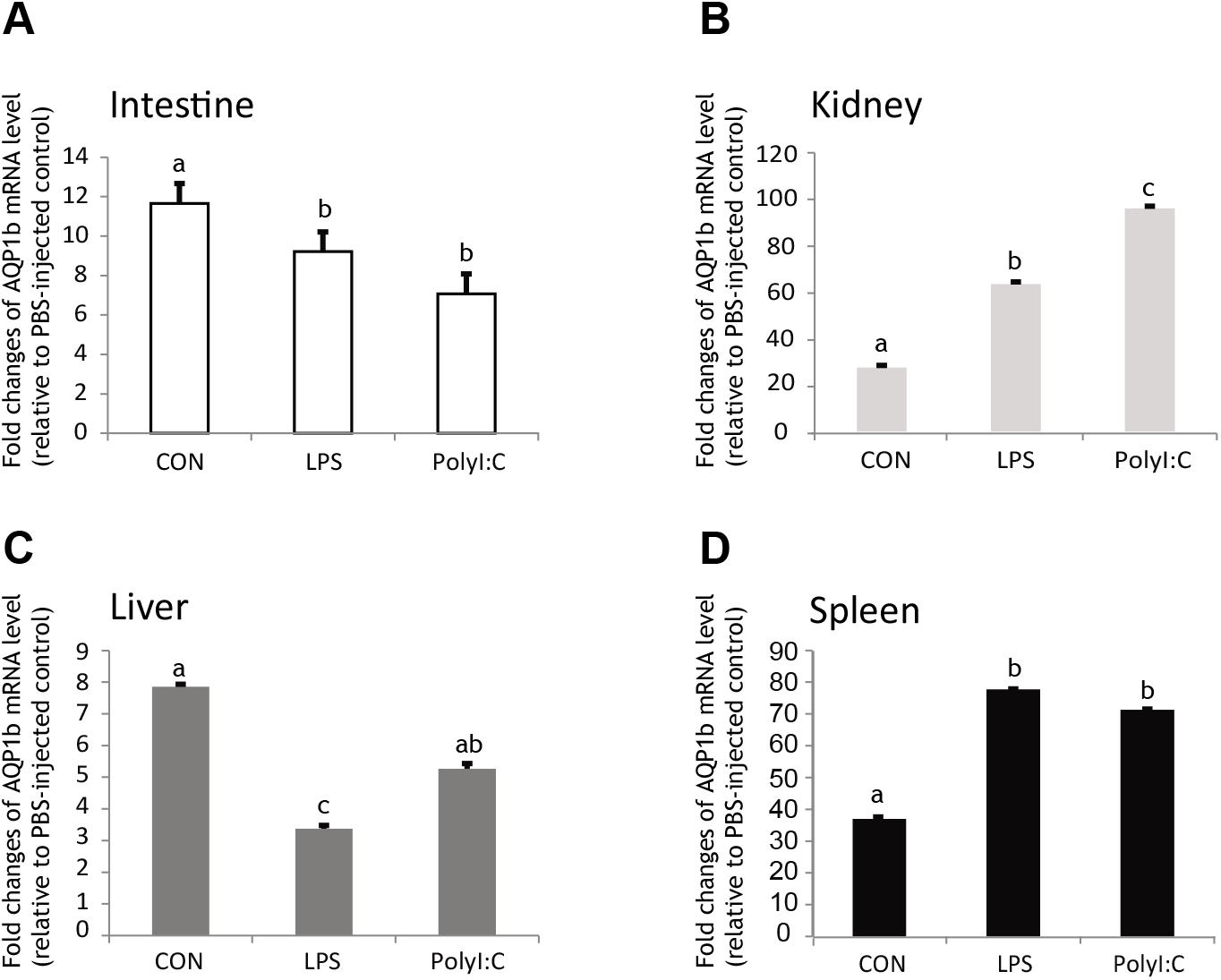
Differential modulation of mmAQP isoforms by immunostimulant exposure. Data represent means ± SDs, with letters indicating significant differences, as analyzed by ANOVA followed by Duncan’s multiple range tests.

## Discussion

We determined that mmAQP1b is similar in sequence and predicted topology to previously identified AQPs. The mud loach AQP1b has traditional structural features of aquaporins, such as six transmembrane domains. These are vital characteristics that appear in the major intrinsic protein (MIP) family and in aquaporins (Borgnia et al., 1999). The tandem repeat structures with two NPA sequences have been proposed to form tight turn structures that interact in the membrane to form the pathway for water to move through the protein (Nielsen et al., 1999). At the genome level, the mmAQP1b gene has a somewhat different organizational structure (*i.e.* 5 exons interrupted by 4 introns) compared to most other teleostean AQP1 orthologues, which have 4 exons (Tingaud-Sequeira et al. 2008, 2010; Kim et al., 2014). We also isolated the complete mRNA of two AQP1b transcript variants. Analysis of mmAQP1b cDNA using available information on the genome organization of the AQP gene in teleost suggest that each isoform is generated by alternative splicing. This leads to a splicing event where the 5’ splice site of the intron is different, which leads to mRNAs with different C-termini. However, the rest of the AQP gene sequence is highly conserved between the two variants.

*In silico* analysis of mmAQP1b promotor identified various putative cis-regulatory elements that may serve targets for sequence-specific transcription factors. We found numerous consensus sequences that may be bound by transcription factors involved in stress and/or innate immunity in teleosts such as STAT, CEBP, CREB, and NFAT5. In particular, NFAT5, a member of the nuclear factor of activated T cell family, plays crucial roles in detecting environmental salinity and immune responses under pathophysiological conditions associated with hyperosmotic stress in teleosts and mammals (Küper et al., 2015; Lorgn et al., 2017). In addition, the canonical motifs for STAT, a key factor in the JAK/STAT pathway were identified, suggesting that AQP1b is involved in inflammation-mediated modulation upon pathogen infection. This observation is consistent with the expression profiles of AQP1a and 3a in kidney, intestine, liver, and spleen of mud loaches after immune challenge (Lee et al., 2017). The mmAQP1b promoter possesses potential CREB sites responsive to cAMP, similar to those found in mammalian and teleost AQP genes (Zapater et al., 2013; Wang and Zheng, 2011). We also observed a potentially conserved site for glucocorticoid-responsive and related elements, which are induced by glucocorticoid or progesterone receptors in mammals and teleosts (Zapater et al., 2013; Lieberman et al., 1993; Moon et al., 1997). Additionally, HNF-1 is a major regulator of glucose homeostasis in the liver, kidney, and pancreas in mammals (Pontoglio, 2000). This evidence indicates that AQP-mediated cellular pathways could be directly or indirectly associated with carbohydrate metabolism in the teleost liver for energy supply during saline challenge.

Interestingly, the *in silico* analysis showed putative binding sites for SOX transcription factors in the mmAQP1b promoter region, similar to several fish species (Cerdà et al., 2013; Zapater, et al., 2013; Wei et al., 2016). Recently, SOX transcription factors were reported to be involved in diverse physiological processes such as the formation of the nervous system (Overton et al., 2002), gonadogenesis (Weiss et al., 2003), or sex determining factor. During embryonic development, mmAQP1b expression was weakly expressed during early embryogenesis, followed by a considerable increase until 2 dph and a subsequent decline. Additionally, AQP1b expression is firstly detected from the onset of fertilization to the 32-cell stage, indicating that AQP1b is maternally inherited, as reported for the common mummicho (*Fundulus heteroclitus*) and zebrafish (Tingaud-Sequeira et al., 2009; Chen et al., 2010). Maternal molecules such as transcripts and protein are provided as a source as of cellular energy, structural components, and defense responses during embryonic and larval development in fish. In addition, there was a decrease in mmAQP1b transcript levels from 4-6 HPF (early blastula-gastrula stage).

In the present study, the expression of mmAQP1b mRNA was detectable in various tissues, including the gill, kidney, and spleen. Some tissues also showed difference in transcript levels. The freshwater teleost kidney is unable to produce hyperosmotic urine, in contrast to seawater-adapted piscine kidneys, which switch to water saving function via expression of divalent ions. Therefore, the function of piscine renal AQP is regulated by environmental salinity (Tipsmark et al., 2010). The expression of AQP1a.2 transcript is not limited to osmoregulatory tissues, but may also occur in non-osmoregulatory tissues (eye, spleen, and testis), as suggested in several teleost species (An et al., 2008; Tingaud-Sequeira, 2010; Kim et al., 2010, 2014; Madsen et al., 2014).

AQP1b transcription showed the significant responses to the immunostimulants LPS and poly(I:C). In particular, renal and splenic AQP1b transcripts were significantly higher than in other tissues. As evidenced by Lehmann et al. (2008), intraperitoneal LPS injection allows LPS immediate access to the circulation. Therefore, LPS induces systemic immune responses. The kidney is the primary excretory organ critical for the maintenance of homeostasis in fish. A recent study suggested that intraperitoneal LPS injection decreases blood flow in the spleen and kidney via vasoconstriction, thereby increasing the volume of interstitial fluid and redistributing blood flow (Wang et al., 2018). Thus, LPS challenge may ultimately impair the transient osmotic gradient, indicating an improper balance of the stable internal water environment and dissolved ion concentrations in the kidney. A previous study reported that upregulated AQP1 transcripts serves a protective role by reversing LPS-induced damage in human renal proximal tubule epithelial cells (Wang et al., 2018). Thus, altered AQP1a protein expression is associated with altered renal physiology.

AQPs have been proposed as molecular osmosensors that maintain water homeostasis. Further, AQPs are a possible regulator of innate host defenses at the level of the plasma membrane (Meli et al., 2018). In the present investigation, we characterized AQP variant transcript levels and investigated AQP1b modulation in response to heavy metal exposure and immune challenge. This study provides a comprehensive basis and strengthens the knowledge of the underlying mechanisms of AQP in physiological and pathological processes. Further studies to deepen the knowledge of fish AQP-mediated mechanisms potentially relevant to molecular pathogenesis are required.

## Competing interests

The authors declare that they have no conflict of interest.

## Funding

This research was supported by the grant from the Korea Institute of Marine Science & Technology (KIMST) funded by the Ministry of Oceans and Fisheries (Project #20170327).

## Authors’ contributions

Sang Yoon Lee contributed to the management of mu loach, gene-cloning, gene-expression analyses and data analysis and; Yi Kyung Kim and Yoon Kwon Nam developed and supervised the experiment, and preparation of the manuscript draft, and modified the manuscript.

## Notes

### Competing Interest Statement

The authors have declared no competing interest.

